# Mitochondria surveillance systems trigger innate immune responses to bacterial pathogens via AMPK pathway in *C. elegans*

**DOI:** 10.1101/2020.09.17.301036

**Authors:** Shouyong Ju, Hanqiao Chen, Shaoying Wang, Jian Lin, Raffi V Aroian, Donghai Peng, Ming Sun

## Abstract

Pathogen recognition and triggering pattern of host innate immune system is critical to understanding pathogen-host interaction. It is generally accepted that the microbial infection can be recognized by host via pattern-triggered immunity (PTI) or effector-triggered immunity (ETI) responses. Recently, non-PRR-mediated cellular surveillance systems have been reported as an important supplement strategy to PTI and ETI responses. However, the mechanism of how surveillance systems sense pathogens and trigger innate immune responses is largely unknown. In the present study, using *Bacillus thuringiensis*-*Caenorhabditis elegans* as a model, we found a new approach for surveillance systems to sense the pathogens through no-PPRs patterns. We reported *C. elegans* can monitor intracellular energy status through the mitochondrial surveillance system to triggered innate immune responses against pathogenic attack via AMP-activated protein kinase (AMPK). Consider that the mitochondria surveillance systems and AMPK are conserved components from worms to mammals, our study suggests that disrupting mitochondrial homeostasis to activate the immune system through AMPK-dependent pathways may widely existing in animals.

## Introduction

Animals encounter diverse pathogens in natural environment and have evolved different defense responses against pathogens for survival. The principal challenge for the host is how to sense the pathogens and trigger defense responses to fight against pathogens. The innate immune system is a universal and evolutionarily ancient part of such host defense response (Janeway & Medzhitov, 2002).

It is generally accepted that hosts could discriminate pathogens from non-pathogenic bacteria in multiple ways. Firstly, the host can recognize microbe-associated molecular patterns (MAMPs) or endogenous danger-associated molecular patterns (DAMPs) by pattern-recognition receptors (PRRs) to induce immune signaling pathways (Takeuchi & Akira, 2010). This is also called pattern-triggered immunity (PTI), which is the most traditional way for hosts identifying pathogens. MAMPs include lipopolysaccharide (LPS), lipoproteins, carbohydrates, flagellin and nucleic acids, which are highly conserved and common among microbes (Wang *et al*, 2020; Yu *et al*, 2017). DAMPs are endogenous molecules secreted by damaged cells, such as interleukin1α, uric acid, S100 cytoplasmic proteins and the extracellular matrix molecule hyaluronan (Van Crombruggen *et al*, 2013). However, the PRR ligands are not unique to pathogenic bacteria, but also found in the non-pathogenic bacteria (Kau *et al*, 2011), making it hard to simply identify the pathogen in this way. Besides, the hosts could sense the existence or the damage caused by certain virulence factors to discriminates pathogens, which is called effector-triggered immunity (ETI) (Bigeard et al, 2015; Galán & Waksman, 2018; Stuart et al, 2013). However, most of the ETI responses still depend on PPRs, while animals could still sense pathogens without activating PPRs (Pukkila-Worley, 2016). Therefore, the PTI and ETI responses relied on PPRs are not enough to explain pathogen recognition (Dangl & Jones, 2001; Dunbar *et al*, 2012).

Recently, there is increasing studies concern about the host could indirectly detect pathogens through non-PPRs-related cellular surveillance responses. In this pattern, pathogens caused severe cell damage to disrupt host core cell processes, while host cellular surveillance systems could monitor such core cellular activities and trigger innate immune responses through non-PPRs-related ways. The change of cellular homeostasis or the core cellular activities caused by pathogens, such as the integrity of nucleolus (Dean *et al*, 2010) or DNA (Ermolaeva *et al*, 2013), inhibit the transcription or translation processes (Dunbar *et al*., 2012; McEwan *et al*, 2012), and disruption the mitochondrial protein folding environment (Liu *et al*, 2014a; Pellegrino *et al*, 2014b), can be sensed by host cellular surveillance systems to initiate innate immune defenses. Consequently, the host cellular surveillance response is an important supplement strategy to PTI and ETI responses during pathogen recognition process. However, despite the continuous enrichment of the immune surveillance systems, the specific upstream signaling molecules and pathways representing the changes of cell homeostasis to induce the innate immune defenses remain largely unknown.

By far, the most classic reported cellular surveillance systems contains the ribosome, proteasome, and mitochondria (Melo & Ruvkun, 2012). The mitochondria are important biosynthetic and bioenergetic organelles and play a key role in cellular energy, metabolism, and protein stability, which also play an important role in the monitor of cell homeostasis (Monlun *et al*, 2017; Roger *et al*, 2017). Numbers of work showed that the mitochondria could be attacked and damaged by toxins produced from pathogens. Toxins from many bacterial pathogens like vacuolating toxin (VacA) (Galmiche & Rassow, 2010), alpha-toxin(Cohen *et al*, 2018), and leucocidin (Genestier *et al*, 2005) can target and damage the mitochondria, leading to mitochondrial dysfunctional. Furthermore, the infection of *P. aeruginosa* caused mitochondrial dysfunctional, while the mitochondrial stress then leads to mitochondrial UPR(UPR^mt^) response by the transcription factor ATFS, which eventually engages host innate immune defenses (Liu *et al*, 2014b; Pellegrino *et al*, 2014a). These studies indicate that the mitochondrial surveillance system is an effective means for host to monitor pathogen infection. However, in addition to sense pathogens through UPR^mt^, other mitochondria mediated ways for host to monitor pathogens and trigger the innate immune responses remains to be studied.

The *Caenorhabditis elegans* has been developed as a powerful model to understand the innate immune responses. Many studies supported that *C. elegans* could monitor core cellular physiology activities to detect pathogen infections (Dunbar *et al*., 2012; McEwan *et al*., 2012; Melo & Ruvkun, 2012). Moreover, there are no exact PPRs that have been unambiguously defined in *C. elegans* (Cohen & Troemel, 2015). These facts make *C. elegans* an ideal model to study how surveillance systems sense the pathogens through a no-PPRs pattern. *Bacillus thuringiensis* (Bt) is an obligate and opportunist pathogen of insect and worms, which produces insecticidal or nematicidal crystal protein during sporulation, and has been used as a leading bio-insecticide to control various pests and worms (Ruan *et al*, 2015). Here, we used a nematicidal Bt strain BMB171/Cry5Ba (Geng *et al*, 2017) and *C. elegans* as a model, to research the detailed mechanisms about how the cell surveillance systems sense pathogens and activate the innate immune responses. We showed that Bt infection causes mitochondrial damage, which ultimately leads to a severely cellular energy imbalance by altered of AMP/ATP ratio. While the Cry toxin-mediated mitochondrial damage will trigger the innate immune response via a cellular energy sensor AMPK in *C. elegans*. Our work revealed the mitochondrial surveillance systems can discriminate pathogens from the non-pathogenic bacteria by triggering innate immune responses via cell energy sensor AMPK during bacteria pathogen infection in *C. elegans*, which provided novel insights to understand host-pathogen interactions.

## Results

### *B. thuringiensis* infection leads to cellular energy imbalance in *C. elegans*

To investigate the intracellular physiological changes of *C. elegans* after pathogenic Bt infection, we conducted the *C. elegans* transcriptome analysis after infection by the nematicidal Bt strain BMB171/Cry5Ba, an acrystalliferous Bt mutant BMB171 transformed with toxin gene *cry5Ba* on the shuttle vector pHT304 (Geng *et al*., 2017). As a control, we compared the transcriptome to a non-nematicidal Bt strain BMB171/pHT304, BMB171 transformed with the empty vector pHT304. Enrichment pathway analyses highlighted several pathways strongly affected by the infection of nematicidal Bt strain (Fig. 1A and Table S2). Interestingly, we found the energy metabolic related pathway was most strongly affected. This indicated that the regulators of the energy metabolic pathway may play important roles in host responses against Bt infection. To confirm this, we measured the concentrations of AMP and ATP by LC-MS (Coulier *et al*, 2006) when wild-type *C. elegans* N2 fed with BMB171/Cry5Ba, the non-nematicidal control strain BMB171/pHT304 and the standard food strain *E. coli* OP50. The results showed that the AMP/ATP ratio had no significant difference after *C. elegans* N2 fed with BMB171/pHT304, but significantly increased after fed with BMB171/Cry5Ba, comparing with that of fed with *E. coli* OP50 (Fig. 1B). However, the AMP/ATP ratio showed no significant difference when the Cry5Ba-receptor mutant *bre-5(ye17)* worms were fed with either BMB171/Cry5Ba or BMB171/pHT304 (Fig. 1B). Next we tested whether other nematicidal Bt can lead cell energy change, such as BMB171/Cry5Ca (Geng *et al*., 2017), BMB171/Cry6Aa (Wei *et al*, 2003), BMB171/Cry21Aa and non-nematicidal BMB171/Cry1Ac (Fang *et al*, 2009). We found that nematicidal Bt strains can cause significant energy imbalance of *C. elegans* comparing with non-nematicidal Bt (Fig. S1). Taking together, we demonstrated that nematicidal Bt infection triggers a cellular energy imbalance of *C. elegans*, which was mainly attributed to the nematicidal toxins.

**Figure 1.**
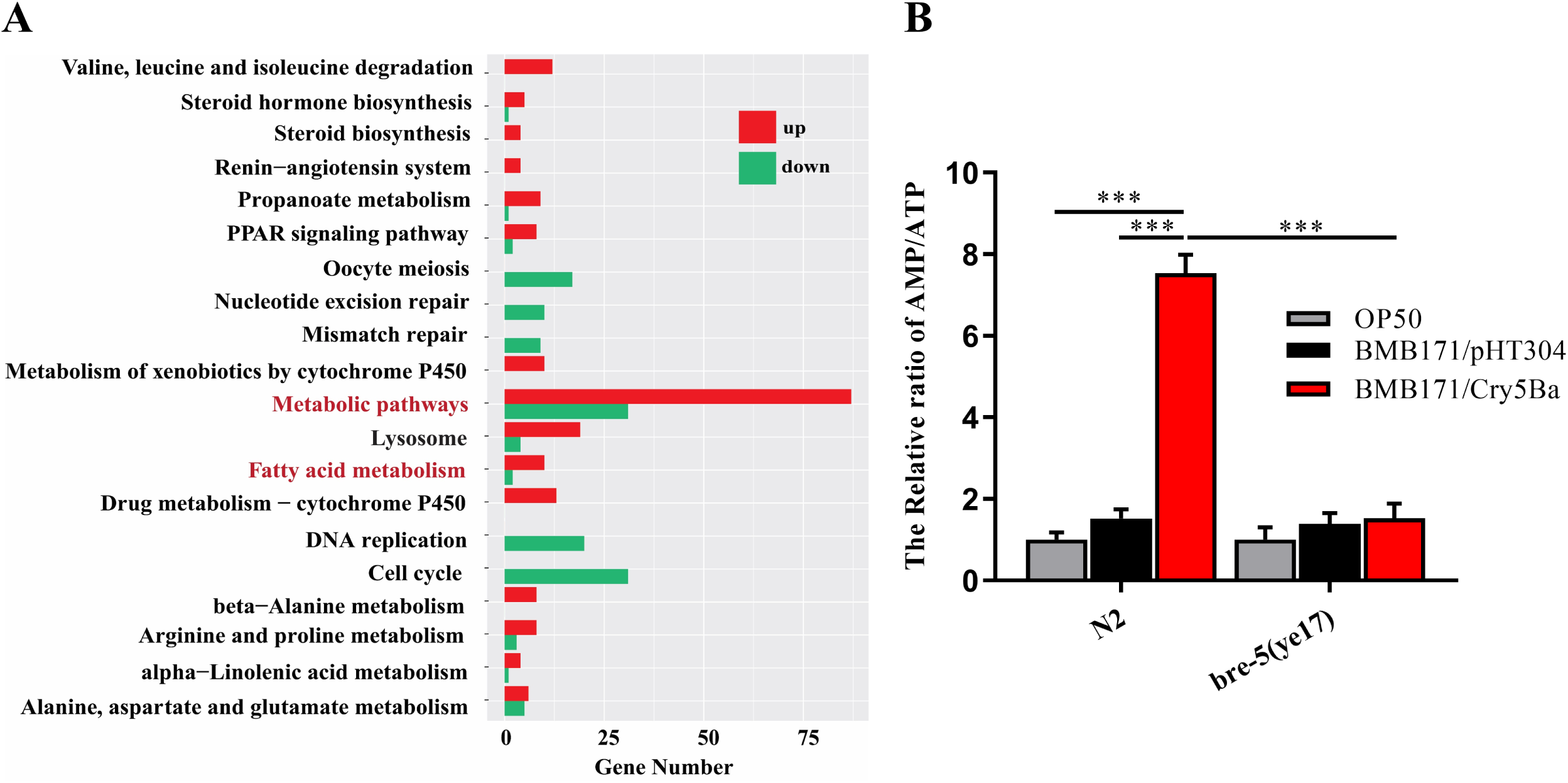
Nematicidal Bt infection leads to the energy imbalance in *C. elegans*. (A) The top 20 pathways affected by the infection of nematicidal Bt obtained by the enrichment pathway analyses of RNA-Seq data. (B) Analysis of the AMP/ATP ratio of wild-type N2 and Cry5Ba-receptor null mutant *bre-5(ye17)* worms fed with OP50, BMB171/pHT304 and BMB171/Cry5Ba. The mean and SD of three independent replicates are shown. Values differences were calculated by one-way ANOVA. Triple asterisks (***) are set at *p*< 0.001, two asterisks (**) are set at *p*< 0.01. a single asterisk (*) is set at *p*<0.05.

### *B. thuringiensis* infection leads to mitochondria damage in *C. elegans*

Mitochondria produces most of the cell’s ATP through oxidative phosphorylation and the tricarboxylic acid cycle, and plays a vital influence for cell metabolism (Rambold & Pearce, 2018). Mitochondria are constantly in a state of fusion and division, and the morphological changes driven by the fusion of mitochondria are critical for maintaining mitochondrial respiration and homeostasis. The dynamic dysfunction of mitochondrial fusion/fission causes mitochondria disruption and cell death (Cogliati *et al*, 2013; Mills *et al*, 2017). It has been reported that the toxins from *Pseudomonas* pathogens can target mitochondria and lead to mitochondria fragmentation (MF) and it dysfunctional (Liu *et al*., 2014a; Pellegrino *et al*., 2014b). The MF phenomenon can result in bioenergetics defects, which cause cell energy imbalance (Liesa & Shirihai, 2013). To assess how nematicidal Bt causes cellular energy imbalance. We tested several physiological and biochemical aspects of mitochondrial damage including MF, mitochondrial membrane potential (ΔΨm), and mitochondrial DNA (mtDNA) content. We used the transgene worms SJ4143(*zcIs17*[P_*ges-1*_::GFP^mt^]) as MF reporter which stably expressed GFP in mitochondria matrix of intestinal cells to detect the mitochondrial morphology (Urano *et al*, 2002). When worms fed with nematicidal Bt BMB171/Cry5Ba, 76.67% of worms showed MF (Fig. 2A and 2B). However, when worms were fed with non-nematicidal Bt strain BMB171/pHT304 or the standard food strain *E. coli* OP50, the mitochondria morphologies of most worms kept tubular, only less than 15% of worms showed MF (Fig.2A and 2B). These results showed that nematicidal Bt infection leads to the MF. Next, we tested whether BMB171/Cry5Ba infection can cause changes in mitochondrial membrane potential (ΔΨm) and mitochondrial DNA (mtDNA) content. Using wild type N2 worms, we found BMB171/Cry5Ba infection can lead to a significant reduction in mitochondrial membrane potential (ΔΨm) and mtDNA content comparing with non-nematicidal Bt BMB171/pHT304 (Fig.2C, 2D and S2).

**Figure 2.**
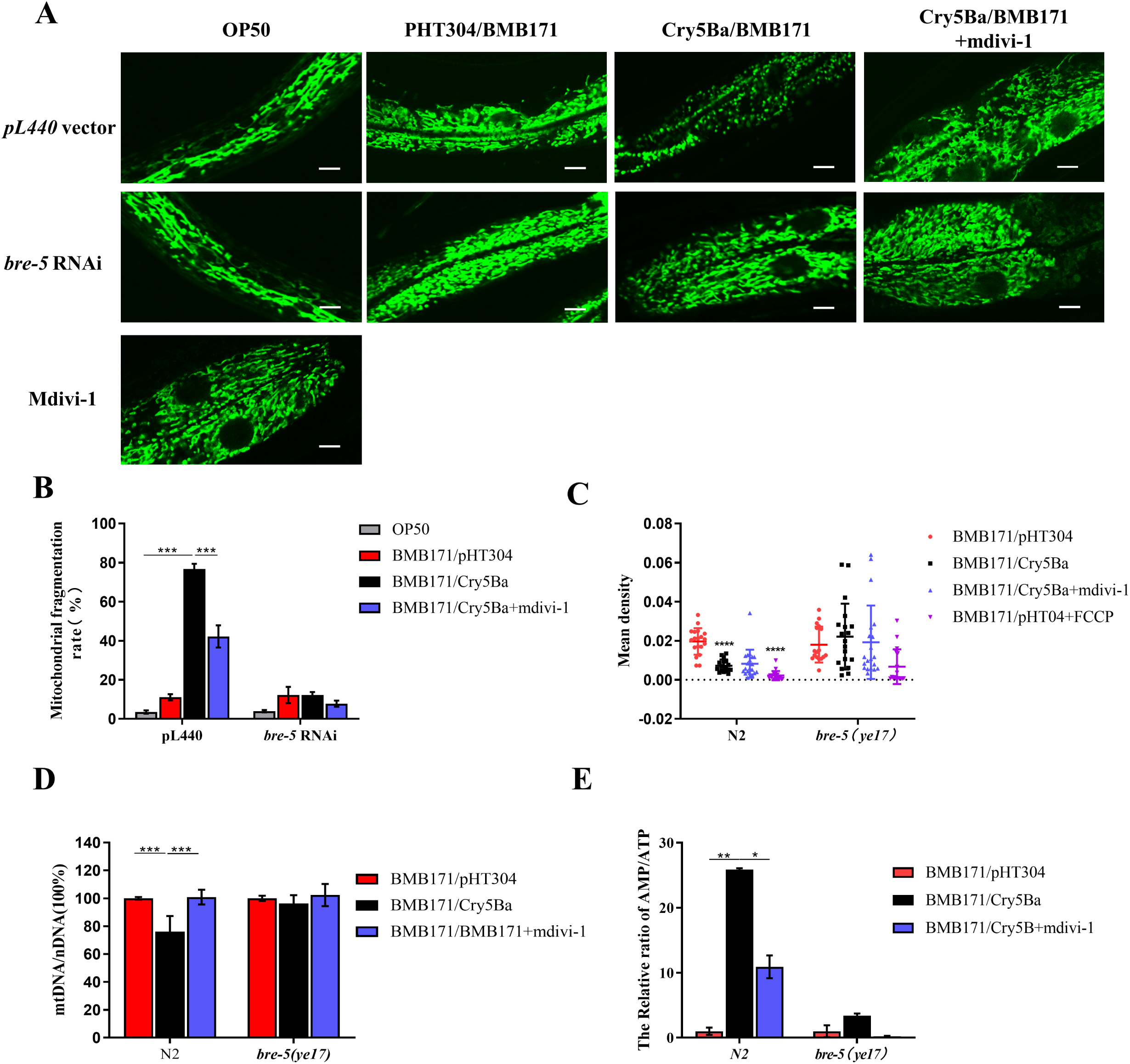
*B. thuringiensis* infection leads to mitochondria damage. (A). Observe the mitochondrial morphology of transgenic *C. elegans* SJ4143(zcIs17 [P_*ges-1*_::GFP^mt^]), which were fed with different bacteria under *bre-5* RNAi or not. The representative images of worms treated by each strain are shown. The scan bars represent 20 μm. (B) Analysis of the percentage of worms that showed MF phenotype after treatment with each Bt strain. Transgenic worms subjected to *bre-5* RNAi or empty vector were treated with different strains.. (C) Analysis of the mitochondrial membrane potential (ΔΨm)) under each treatment. The level of ΔΨm was determined by densitometry of each worm. Around 20 worms were measured in each condition. (D) The relative ratio of mtDNA /nDNA was investigated by Real-time PCR and then analyzed. (E) The relative AMP/ATP ratio was measured in each condition. In each assay, the mean and SD of three independent replicates are shown. Values differences were calculated by one-way ANOVA. Triple asterisks (***) are set at *p*< 0.001, two asterisks (**) are set at *p*< 0.01. a single asterisk (*) is set at *p*<0.05.

When the Cry5Ba receptor related gene *bre-5* was silenced by RNA interference (RNAi) in transgene worms SJ4143(*zcIs17* [P_*ges-1*_:: GFPmt]), only 12.22% of worms showed MF phenomenon after fed with BMB171/Cry5Ba (Fig. 2A and 2B), indicating that the Cry protein toxin Cry5Ba was the key factor that caused the MF phenomenon during Bt infection. However, when Cry5Ba-receptor null mutant *bre-5*(*ye17*) worms fed with BMB171/Cry5Ba in the same condition, the ΔΨm and mtDNA content were similar to *bre-5*(*ye17*) worms fed with BMB171/pHT304 (Fig 2C-D). We conclude that Cry5Ba toxin is the key factor for Bt to manipulate host cell mitochondria and cause serious mitochondrial dysfunction.

Mdivi-1 is an efficient inhibitor to attenuates mitochondrial division by inhibiting the mitochondrial division dynamin, and it also suppresses mitochondrial outer membrane permeabilization (Cassidy-Stone *et al*, 2008). To assess the relationship between cell energy imbalance and the mitochondrial dysfunction after Bt infection, we checked whether mitochondrial division inhibitor mdivi-1 can recover mitochondria damage. We observed that mdivi-1 can effectively reduce Cry5B-mediated MF (Fig. 2A-B). Our results showed that when wild type N2 worms were fed with BMB171/Cry5Ba adding into mdivi-1, the AMP/ATP ratio was significantly reduced, the ratio of mtDNA/nDNA returned to normal levels (Fig. 2D-E). However, we found mdivi-1 cannot recover the reduction of mitochondrial membrane potential (Fig. 2C). In general, these results indicate that mitochondrial damage is responsible for the increase in intracellular AMP/ATP.

To assess whether other nematicidal Bt strains can cause MF phenomenon, we fed transgene worms SJ4143(z*cIs17* [P_ges-1_:: GFP^mt^]) with the other four nematicidal Bt BMB171/Cry5Ca, BMB171/Cry21Aa, BMB171/Cry6Aa, and non-nematicidal Bt BMB171/Cry1Ac. These former two strains produce Cry5-like three-domain (3D) group nematicidal Cry proteins Cry5Ca (Geng *et al*., 2017) and Cry21Aa (Pardo-Lopez *et al*, 2013), respectively. While BMB171/Cry6Aa produces a non-3D group nematicidal Cry toxin Cry6Aa (Zhang *et al*, 2016). The transgene worms SJ4143(z*cIs17* [P_ges-1_:: GFP^mt^]) fed with nematicidal BMB171/Cry5Ca and BMB171/Cry21Aa exhibited the fragmented mitochondrial morphology (Fig. S3) with a range from 65.42% to 86.73% (Fig. S4). However, the worms fed with the BMB171/Cry6Aa and BMB171/Cry1Ac showed no significant MF phenomenon (Fig. S3 and Fig. S4). The results showed that the nematicidal Bt Cry5-like toxins are capable of causing the mitochondrial morphology damage in *C. elegans*.

### Cell energy imbalance mediated by mitochondria damage activates the AMP-activated protein kinase

The AMP-activated protein kinase (AMPK) is a sensor of energy status that maintains energy homeostasis and can be activated by a decrease in energy levels (Lobet *et al*, 2015). At this low energy level, it switches on catabolic pathways that generate ATP, while switches off ATP-consuming anabolic processes (Lobet *et al*., 2015). AMPK is activated via phosphorylation of Thr172 on the α catalysis subunit (AAK-2 protein) (Lobet *et al*., 2015). It is well known that increasing of AMP/ATP proportion is the classical way to activate the AMPK (Lobet *et al*., 2015). We observed that *aak-2* was significantly up-regulated after Bt infection (Fig. S5), and our results showed that AMP/ATP ratios are significantly increased when worms are infected by nematicidal Bt (Fig. 1B). Therefore, we speculated that nematicidal Bt infection may activate AMPK. To confirm this hypothesis, we performed western blotting to analyze the Thr172 phosphorylation of AAK-2 protein in worms. The results showed that the Thr172 of AAK-2 protein was phosphorylated when *C. elegans* fed with BMB171/Cry5Ba but not in the control strain BMB171/pHT304 treatment (Fig. 3A). What’s more, the Thr172 of AAK-2 protein was not phosphorylated when Cry5Ba-receptor null mutant *bre-5*(*ye17*) worms were fed with BMB171/Cry5Ba under the same conditions (Fig. 3A). Adding mdivi-1 could significantly suppress the Thr172 phosphorylation of AAK-2 (Fig. 3A). These results demonstrated that AMPK of worms is activated by nematicidal Bt infection via the phosphorylation of the core subunit AAK-2.

**Figure 3.**
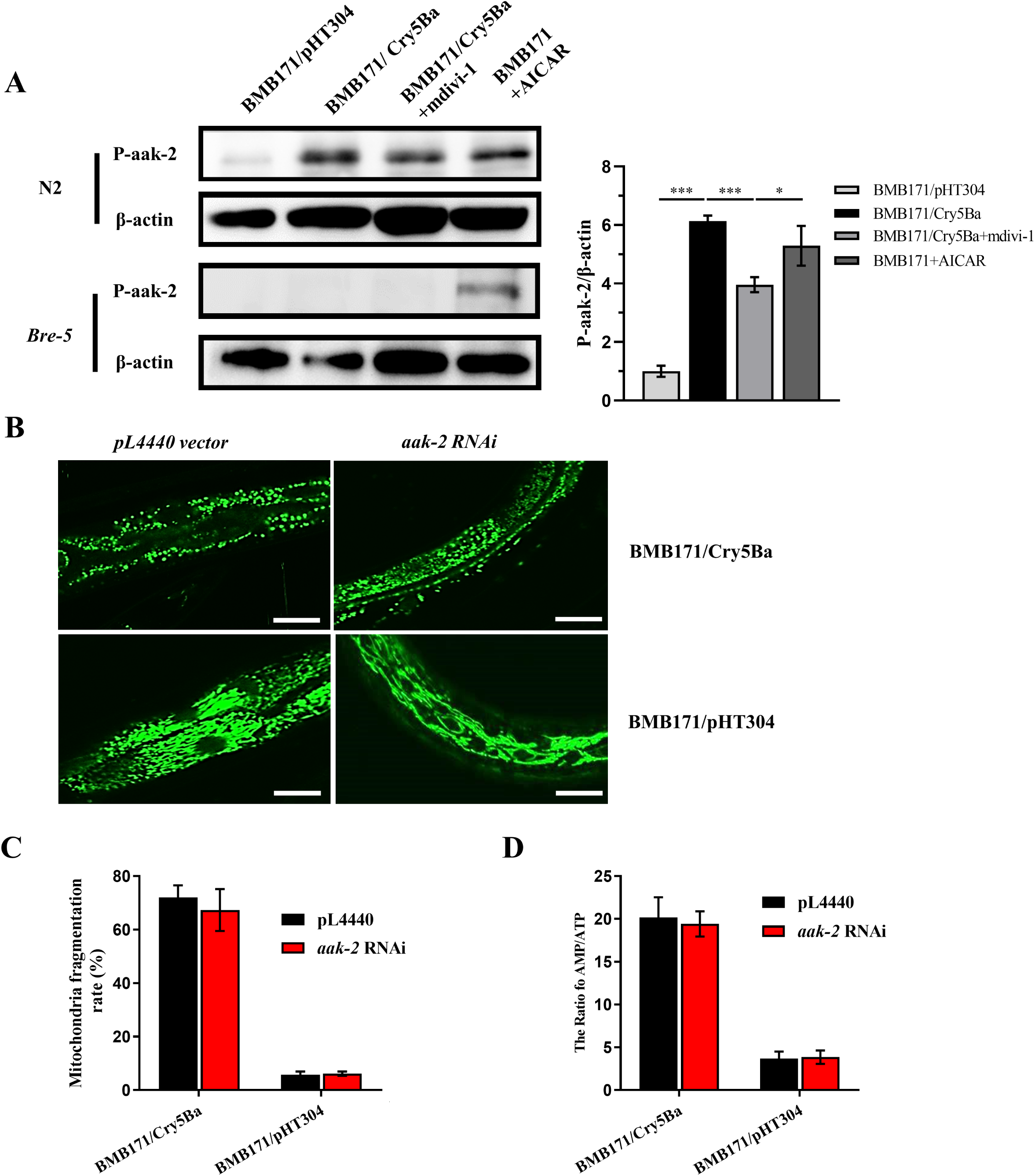
AMPK was induced by nematicidal Bt infection. (A) Left: western blotting showed that AMPK was activated by the phosphorylation of the protein AAK-2 when exposed to BMB171/Cry5Ba after 4 h. Right: amounts of p-AAK-2 were determined by densitometry of protein bands from three independent experiments. β-actin was the loading control. (B) The mitochondria morphologies observations of transgenic *C. elegans* SJ4143(zcIs17[P_*ges-1*_::GFP^mt^]) fed with BMB171/Cry5Ba or BMB171/pHT304. Representative images of worms treated by each strain are shown. The scan bars represent 20 μm. (C) After *aak-2* RNAi, analysis of the percentage of worms showing MF phenotype after treatment with BMB171/PHT304 and BMB171/Cry5Ba. (D) Analysis of the AMP/ATP ratio after treated by BMB171/pHT304 and BMB171/Cry5Ba when *aak-2* RNAi. In each condition, the mean and SD of three independent replicates are shown. Values differences were calculated by one-way ANOVA. Triple asterisks (***) are set at *p*< 0.001, a single asterisk (*) is set at *p*<0.05.

To assess the relationship between cell energy imbalance and the activating of AMPK after Bt infection, we knock down the *aak-2* transcription levels by RNAi in the mitochondria reporter worms SJ4143(zcIs17[P_ges-1_:: GFP^mt^]). Then, the non-RNAi and *aak-2* RNAi worms were fed with BMB171/Cry5Ba or BMB171/pHT304, respectively. The RNAi-*aak-2* worms also showed a high level of MF when worms fed with BMB171/Cry5Ba after 5 h, most mitochondria keep tubular when worms fed with control strain BMB171/pHT304 (Fig. 3B).

We also measured the concentrations of AMP and ATP when wild-type *C. elegans* N2 and RNAi-*aak-2* worms were fed with BMB171/Cry5Ba or BMB171/pHT304. The AMP/ATP ratio was significantly increased after RNAi-*aak-2* worms were treated with BMB171/Cry5Ba compared to worms fed with the control strain BMB171/pHT304 (Fig. 3C). However, the increasing of AMP/ATP ratio was no significant difference when RNAi-*aak-2* and N2 worms were treated with BMB171/Cry5Ba (Fig. 3D). These results demonstrate that knockdown of the *aak-2* does not affect the level of MF and energy imbalance during Bt infection, and suggest that AMPK activation is a result rather than a cause of the above MF phenomenon. Thus, we concluded that Bt-mediated cell energy imbalance activates the AMPK in *C. elegans*.

### AMPK activation is involved in *C. elegans* defense responses against Bt infection

Several previous studies reported that AMPK defends against low glucose levels, dietary deprivation, paraquat, physical stress and pathogens (Brunton *et al*, 2013; Fukuyama *et al*, 2012; Herzig & Shaw, 2018). Therefore, we asked whether AMPK activation might be involved in defense against Bt infection. To confirm this hypothesis, the growth of different AMPK deficient mutant worms were quantitated relative to that of wild type N2 worms when fed different doses of spores and toxin mixtures from BMB171/Cry5Ba (Bischof *et al*, 2006). There are three subunits for the AMPK complexes, including a catalytic subunit (α), and two regulatory subunits (β and γ). We tested the sensitivity of the four AMPK null alleles mutants and the wild type N2 worms exposed to BMB171/Cry5Ba, including *aak-1*(*tm1944*) (subunit α1 of AMPK), *aak-2*(*ok524*) (subunit α2 of AMPK), *aakb-1*(*tm2658*) (subunit β1 of AMPK), and *aakg-4*(*tm5269*) (subunit γ1 of AMPK), etc. Compared to the wild type N2 worms, only *aak-2*(*ok524*) mutant worms showed more increased sensitivity to BMB171/Cry5Ba infection (Fig. 4A and Fig. 4B). Survival assays confirmed that the *aak-2*(*ok524*) mutant worms are more sensitive to BMB171/Cry5Ba infection (Fig. 4C).

**Figure 4.**
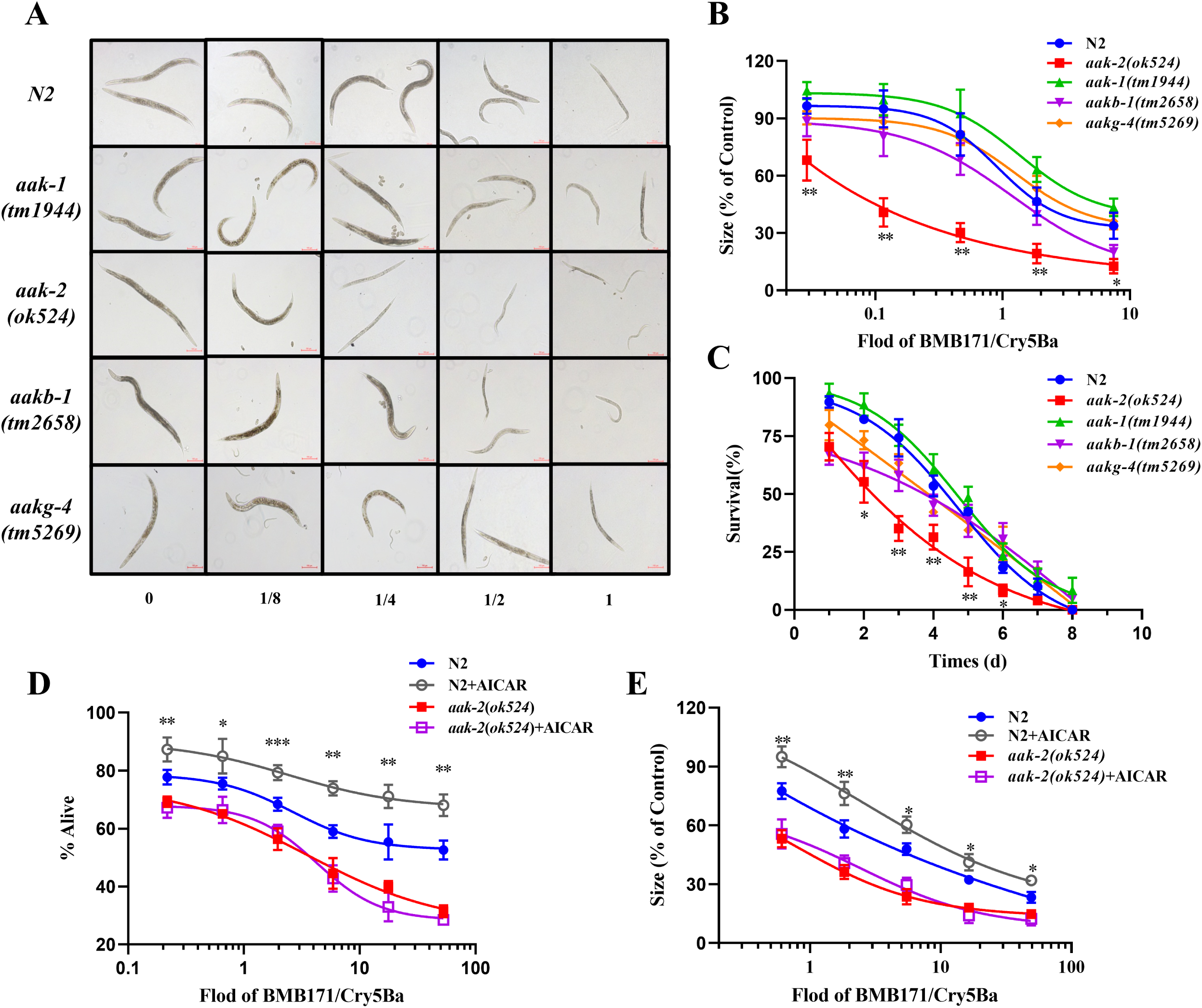
The AMPK activation is involved in C. elegans defense responses against Bt infection. (A) The growth assay of the wild type N2 and mutant worms after BMB171/Cry5Ba infection. The representative worms are shown for each dose. Scale bar, 0.02 mm. (B) To photographed at least 60 worms for each dose under 100 magnification microscope, calculated the average area for each condition. The size of the worms in the absence of toxin was set at 100%. (C) The survival assay of the wild type worms N2 and mutant worms after BMB171/Cry5Ba infection. The survival assay (D) and the growth assay (E) of the wild type worms N2 and mutant worms *aak-2*(*ok524*) after BMB171/Cry5Ba infection when the AMPK was activated by AICAR or not. The mean and SD of three independent replicates are shown. Asterisks indicate significance as determined by a Student’s t-test (*, *p*<0.05; **, *p*<0.01) and ns indicate no significant difference.

To further confirm the importance of AMPK catalysis subunit α2 (AAK-2 protein) in defense to BMB171/Cry5Ba infection, we tested by the growth assay the sensitivity to another null allele of *aak-2* mutant, *aak-2*(*gt33*) feeding on BMB171/Cry5Ba. We found that *aak-2*(*gt33*) is also more sensitive to BMB171/Cry5Ba than wild type N2 (Fig. S6). A similar phenotype was observed in *aak-2* RNAi worms (Fig. S7).

We next asked whether the AMPK subunit α2 knock out phenotype is specific for Bt infection. The sensitivity of the wild type N2 and *aak-2*(*ok524*) mutant worms were analyzed to the heavy metal copper sulfate and oxidative stress (hydrogen peroxide). Our results showed that it is no significant difference in these treatments between the wild type N2 and the mutant *aak-2*(*ok524*) worms (*t*-test, *p*> 0.05) (Fig. S8A and S8B).

In addition, we activated AMPK using 5-Aminoimidazole-4-carboxamide 1-ß-D-ribofuranoside (AICAR), a typical activator stimulating AAK-2 kinase activity of AMPK via phosphorylation of Thr172 on α catalysis subunit (AAK-2 protein) (Kristiansen *et al*, 2009). Western blotting results showed that the AMPK is activated by 50 µM of AICAR via AAK-2 Thr172 phosphorylation (Fig. 3A). We then compared the sensitivity of the AICAR-treated N2 worms with non-treated N2 worms to BMB171/Cry5Ba infection by survival assays and growth assay, respectively. The results showed that the AICAR-activating worms showed more significant resistance to BMB171/Cry5Ba infection comparing with the no-activating N2 worms (Fig. 4D and 4E). However, compared with the wild type N2 worms, the resistance to BMB171/Cry5Ba infection was not significant difference for *aak-2* mutant either in AICAR-activating or no-activating conditions (Fig. 4D and 4E). Taking together, our results demonstrated that AMPK plays important roles for *C. elegans* resistance to attack by BMB171/Cry5Ba.

### AMPK activity in the intestine is required for *C. elegans* resistance to Bt infection

To independently confirm the importance of AAK-2 in defense to BMB171/Cry5Ba infection, we constructed the transgenic strain *aak-2*(*ok524*) (P_*aak-2*_::*aak-2*), which expresses the *aak-2* under the control of its native promoter P_*aak-2*_ to rescue the *aak-2* function under the background of mutant *aak-2*(*ok524*) worms. The growth assay and survival assay results showed that *aak-2* expression driven by its own promoter P_*aak-2*_ can completely alleviate the hypersensitivity of *aak-2*(*ok524*) mutant to BMB171/Cry5Ba infection (Fig. 5A and 5B), supporting that AAK-2 is independently importance for worms to defense BMB171/Cry5Ba infection.

**Figure 5.**
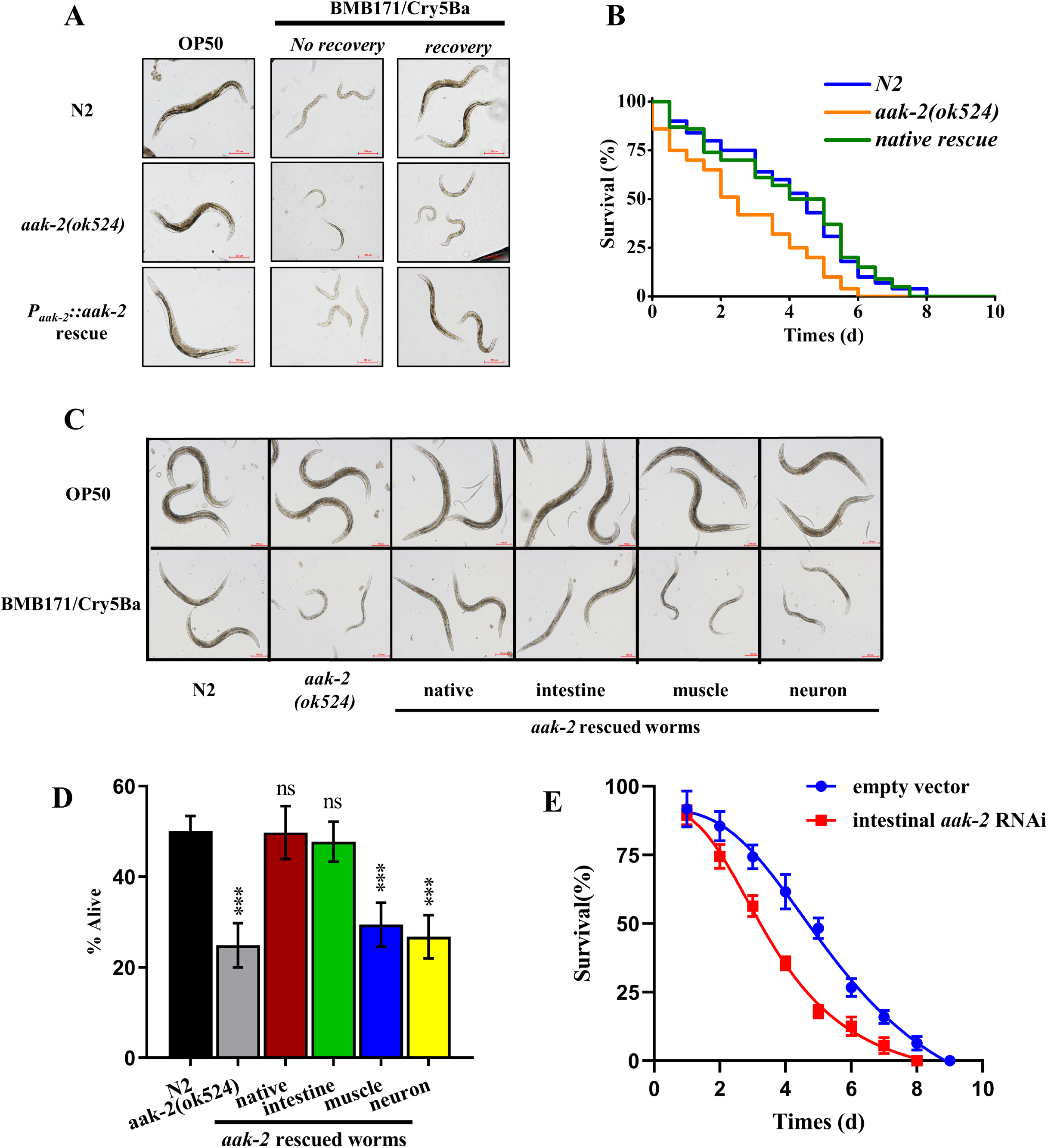
AMPK activity in the intestine is required for *C. elegans* resistance to Bt infection. **(A)** The growth assay of the *C. elegans* N2, *aak-2(ok524)* and P_*aak-2*_::*aak-2* rescue worms after fed with OP50 or BMB171/Cry5Ba for 30 min and then recovery in *E. coli* OP50 NGM plate for 24h or not. The representative images of worms in each condition are shown. (B) The survival assay of the *C. elegans* N2, *aak-2*(*ok524*) and P_*aak-2*_::*aak-2* rescue worms after treated by BMB171/Cry5Ba. (C) The growth assay of the *C. elegans* N2, *aak-2(ok524)* and several kinds of rescue worms after fed with OP50 or BMB171/Cry5Ba. The representative images of worms in each condition are shown. (D) The survival assay of the wild type worms N2, *aak-2(ok524)* and several kinds of rescue worms after BMB171/Cry5Ba infection. (E) The survival assay of the N2 worms after BMB171/Cry5Ba infection when intestinal *aak-2* RNAi or not. The mean and SD of three independent replicates are shown. Values differences were calculated by one-way ANOVA. Triple asterisks (***) are set at *p*< 0.001 and ns indicates no significant difference.

*C. elegans* apparently lacks professional immune cells, but can rely on epithelial cells for immune defenses (Kuo *et al*, 2018). Bt Cry5B mainly acts by attacking the intestine of worms and forming pores in the membrane of the intestine cell (Griffitts & Aroian, 2005; Huffman *et al*, 2004). We therefore hypothesized intestinal-specific activity of AMPK regulates immune responses to Bt infection. We drove the *aak-2* expression under different tissue-specific promoters, including the intestine-specific promoter P_*vha-6*_ (Oka *et al*, 2001), the muscle-specific promoter P_*myo-3*_ (Okkema *et al*, 1993), and the neuron-specific promoter P_*rab-3*_ (Nonet *et al*, 1997). We found that only *aak-2* expression under the intestine-specific promoter P_*vha-6*_ alleviated the hypersensitivity of *aak-2*(*ok524*) mutant to BMB171/Cry5Ba infection (Fig. 5C and 5D). In contrast, there was no significant difference in hypersensitivity to Bt infection among the *aak-2*(*ok524*) mutant and the muscle-specific P_*myo-3*_ or neuron-specific P_*rab-3*_ rescued worms (Fig. 5C and 5D). In addition, we knocked down *aak-2* gene by RNAi in the intestine, muscle, and epidermis of worms, respectively. Only the intestine-specific knock-down of *aak-2* made the worms more sensitive to BMB171/Cry5Ba infection (Fig. 5E). In contrast, epidermal-specific or muscular-specific *aak-2* RNAi worms did not (Fig. S9A and S9B). These results suggest that the intestine serves as the first line of AMPK-mediated defense against BMB171/Cry5Ba attack.

### AMPK activation triggers DAF-16 dependent innate immune singling pathway during Bt infection

AMPK pathway is an evolutionarily conserved from *C. elegans* to mammals and regulates many downstream pathways (Hardie, 2011). Here, AMPK has been shown to play a role in defense of Bt infection in *C. elegans*. However, the AMPK downstream genes or pathways involved in the defense against Bt infection in *C. elegans* were not clear. It was reported that AAK-2 (AMPK subunit α2) is capable of modulating the phosphorylation of the FOXO family transcription factor DAF-16, activating the DAF-16-dependent transcription when worms suffering from dietary restriction (Greer *et al*, 2007). DAF-16 itself regulates many genes, involved in metabolism, immune responses against several pathogens, and longevity of *C. elegans* (Murphy *et al*, 2003; Zečić & Braeckman, 2020). Moreover, the DAF-16 was also triggered by nematicidal Bt infection (Wang *et al*, 2012), and functioned as an important modulator in defense against nematicidal PFTs in *C. elegans* (Chen *et al*, 2010). Therefore, we speculated that AMPK may regulate the DAF-16-dependent signaling pathway in defense against Bt infection. To confirm this hypothesis, we compared previously identified DAF-16 target genes (Murphy *et al*., 2003) with our RNA-Seq data. We found 60 of the genes up-regulated by Bt infection are also the targets of DAF-16 (Fig. 6A, Table S3 and S4) (*t*-test, *p* < 0.001). To verify this analysis, we selected 8 typical genes from these genes and determined their transcription by qPCR. The transcription of these 8 genes was significantly up-regulated after Bt infection in wild type N2 worms. Moreover, RNAi of *daf-16* significantly suppressed the up-regulation of these genes induced by Bt infection (Fig. 6B), and RNAi of *aak-2* also suppressed most gene upregulation, supported that these of DAF-16-dependent genes are also regulated by AAK-2 during Bt infection.

**Figure 6.**
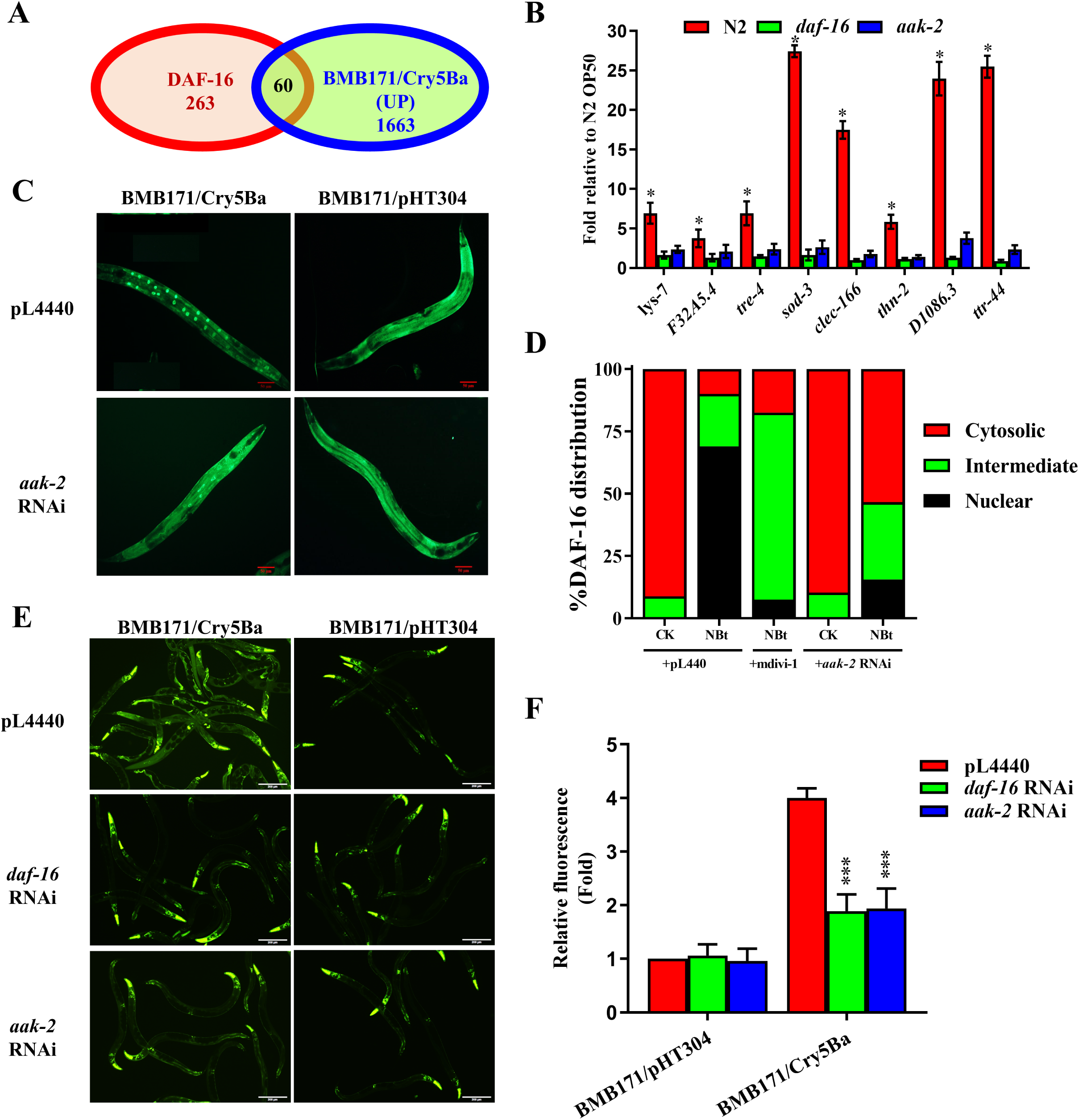
AMPK is required for DAF-16 nuclear accumulation during Bt infection. (A) Venn diagrams comparing show the overlaps in genes activated by nematicidal Bt and the target genes of DAF-16 in C. elegans. (B) qPCR analysis of expression of DAF-16 target genes in wild-type N2, *aak-2*(*ok524*) and *daf-16*(*mu861*) mutant worms after Bt infection. (C) DAF-16 translocation was observed during the no-RNAi and *aak-2* RNAi transgenic worms TJ356(*Isdaf-16*::*gfp*) fed with BMB171/Cry5Ba or BMB171/pHT304 for 2 h, respectively. The representative images for each treatment are shown. Scale bar, 20 µm. (D) The quantification of DAF-16 distribution after Bt infection (CK: BMB171/pHT304; NBt: BMB171/Cry5Ba). The worms with DAF-16 locations as cytosolic (Cyt), intermediate (Int), or nuclear (Nuc) were counted, and the percentages of each pattern of DAF-16 nuclear translocation are calculated. (E) The expression level of P_*sod*_::GFP were observed using a reporter strain CF1553 (P_*sod*_::GFP) during the *aak-2* RNAi, daf-16 RNAi or not after Bt infection. Scale bar, 20 µm. (F) The quantification of P_*sod*_::GFP expression levels calculated by the fluorescence intensity. The mean and SD of three independent replicates are shown. Values differences were calculated by one-way ANOVA. Triple asterisks (***) are set at *p* <0.001, two asterisks (**) are set at *p* <0.01, one asterisk was set at p < 0.05 and ns indicate no significant difference.

Under standard growth conditions, DAF-16 is distributed predominately throughout the cytoplasm of all tissues. When activated, DAF-16 will be phosphorylated and translocated from cytoplasmic to the nucleus, then binds to the promoter region and activates the expression of target genes (Murphy *et al*., 2003). Therefore, we monitored the cellular translocation of DAF-16 using a transgenic worms TJ356(*Isdaf-16*:: *gfp*) as a reporter, which expresses functional DAF-16:: GFP fusion protein. The reporter worms were fed with BMB171/Cry5Ba and a control strain BMB171/pHT304 for 2 h then observed under the fluorescence microscope. After BMB171/Cry5Ba infection, most of the DAF-16:: GFP was translocated from the cytoplasm to the nucleus in the intestine, especially the front and middle parts of the intestines (Fig. 6C and 6D). In contrast, the control strain BMB171/pHT304 failed to cause DAF-16 translocation to the nucleus at the same conditions (Fig. 6C, 6D and Fig. S12). These results demonstrated that nematicidal Bt infection can activate the DAF-16 nuclear translocation in wild-type worms.

To test whether the AMPK activity is required for the activation of DAF-16 during Bt infection, we monitored the cellular translocation of DAF-16 when the *aak-2* gene was silenced by RNAi in the reporter worm TJ356 (*Isdaf-16*::*gfp*). The observations showed that the DAF-16 nuclear translocation induced by Bt is significantly diminished by RNAi *aak-2* gene (Fig. 6C and 6D). To test whether DAF-16 activation can trigger the transcription of the downstream immune-related effectors, we tested the expression of *sod-3*, a well-known direct target of DAF-16 during Bt infection (Hsu *et al*, 2003); *sod-3* gene is also the one of the most significant up-regulated genes in Bt-infected worms (Table S3 and S4). We observed the expression of *sod-3* using transgenic worm CF1553(*muIs84*(sod-3::GFP)) (Hsu *et al*., 2003). When the SOD-3::GFP reporter worms were exposed to BMB171/Cry5Ba for 2 h, the expression of *sod-3* was significantly up-regulated compared with the worms exposed to the control stain BMB171/pHT-304 for 2 h. Meanwhile, the induction of *sod-3* by BMB171/Cry5Ba infection was significantly inhibited when either *daf-16* or *aak-2* gene was silenced in the strain CF1553(*muIs84*(sod-3::GFP) (Fig. 6E and 6F). Additionally, the *aak-2* deletion also significantly suppressed the up-regulation of the above selected 8 immune-related genes induced by Bt infection (Fig. 6B). Taking together, we concluded that the AMPK triggers the DAF-16-dependent innate immune pathway during Bt infection.

## Discussion

Pathogen recognition and triggering the host innate immune system is critical to understanding pathogen-host interaction (Zhang & Zhang, 2009). It is generally accepted that the microbial infection can be recognized by host PTI and ETI patterns. PTI response is related to PRRs, which identify conserved microbial molecules or signal molecules leaked from damaged cells. However, due to the limitations of PRR ligands, PTI is not enough to distinguish pathogens, *e*.*g*., from some probiotic or symbiotic bacteria. The ETI hypothesis is an important supplement to PTI and proposes that the host responds to pathogens by monitoring bacterial virulence factors (Kanyuka & Rudd, 2019; Stuart *et al*., 2013). Moreover, cellular surveillance systems, including the ribosome, proteasome, and mitochondria, monitor core cellular physiology activities were considered as a novel pattern for hosts to distinguish pathogens from other microorganisms (Melo & Ruvkun, 2012). However, how host cells using these systems to sense pathogens are unclear. There are several ways for the host to recognize the pathogens by monitoring core cell processes, including the DNA damage (Ermolaeva *et al*., 2013), the inhibition of translation (McEwan *et al*., 2012), and the UPR^mt^ during the mitochondrial damage state (Melber & Haynes, 2018), etc. Here, we revealed that host cells could sense pathogens via mitochondria mediated intracellular energy imbalance. To our knowledge, there was no report showing host cells can monitor pathogens through energy changes yet. Therefore, our work provides a novel insight for the host cell to detect pathogens through cellular surveillance systems.

Previous works have shown several pathogens could attack and disrupt host mitochondria (Osborne & Brumell, 2017; Pellegrino *et al*., 2014b; Stavru *et al*, 2011). In our study, damage to the mitochondria was also detected when worms were infected by nematicidal Bt, including MF, a decrease in mitochondrial membrane potential, and the changes in mtDNA content (Fig. 2). The damages to the mitochondria during Bt infection was mainly caused by its key virulent factor, Cry5Ba toxin (Fig. 2 and S2). Although Cry5Ba could be translocated into epithelial cells, we did not find evidence of colocalization with mitochondria (Fig. S10), indicating Cry5Ba may lead to mitochondrial dysfunction indirectly. Although the detailed mechanisms of how Cry5B caused mitochondrial damage remain unknown, our data suggests the action model of mitochondrial network disruption by Cry5Ba toxin is likely different from the action model of toxins produced by *P. aeruginosa* (Pellegrino *et al*., 2014b) or *L. monocytogenes* (Stavru *et al*., 2011).

Cellular energy imbalance is an important sign of mitochondrial disorder. In fact, several PFTs produced by pathogens, such as streptolysin O, *Vibrio cholera* cytolysin, *Staphylococcus aureus* α-toxin, and *Escherichia coli* hemolysin A, could also cause a decrease in intracellular ATP levels in a non-virally transformed human keratinocyte cell line (Kloft *et al*, 2010). Our results showed that Bt infection could cause a Cry-dependent cellular energy imbalance, which was widespread among nematicidal Bt infection (Fig. 1B and S1). Cry5Ba is a PFTs that can insert into the membrane of *C. elegans* epithelial cells to form pores in the membrane (Aroian & van der Goot, 2007), which may induce the intracellular substances leakage and cause mitochondrial stress or decrease the concentration of intracellular ATP directly. Both mitochondrial dysfunction and cytosolic energy imbalances were restored in a Cry5Ba receptor mutant (*bre-5*), indicating Cry5Ba is critical for these phenomena. Conversely, the mitochondrial division inhibitor mdivi-1, which can inhibit mitochondrial fragmentation but not the formation of toxin pore, can protect cells from changes in AMP/ATP ratio. Thus, the pores caused by the toxin are not the only reason for the changes in intracellular energy, and mitochondrial are also involved. Overall, these results strongly suggest that the Cry5Ba-like toxin is mainly responsible for cellular energy imbalance during nematicidal Bt infection. Moreover, the Bt strains that produce Cry5B-like nematicidal Cry toxins are capable of disrupting host mitochondria (Fig. S3 and Fig. S4) but not the others. This data indicates that pathogens could cause host energy imbalances in different ways. Of course, the detailed ways and mechanisms of such process still need further research.

In the past few years, accumulating evidence suggests that AMPK is not only a crucial evolutionarily conserved cellular energy sensor but also plays an important role in host-pathogen interactions (Brunton *et al*., 2013). Previous research indicated that cytosolic energy imbalance and calcium content are the important AMPK triggering patterns (Carling *et al*, 2011). By measuring cytoplasmic fluorescence using the calcium indicator Fluo-4AM (Zhang *et al*., 2016), we found Bt infection did not induce the increase of cytoplasm calcium level of worms (Fig. S11A and S11B). Rather, our results indicated that nematicidal Bt infection lead to mitochondria damage, which then gave rise to the dramatic increase of AMP/ATP ratio and AMPK activation via phosphorylation of α-catalysis subunit protein AAK-2 (Fig. 2A). Our data suggest that the increase of AMP/ATP rate activates AMPK, but not the Ca^2+^ content, during Bt infection.

To explain how the cellular mitochondria surveillance system activates AMPK through energy disorder. We found that the inhibition of mitochondrial MF phenotype by the mitochondrial division inhibitor mdivi-1 recovered the mtDNA content, restored abnormal AMP/ATP content, and inhibited the activation of AMPK. These data suggest that MF may be the reason why the AMP/ATP content changed when the mitochondria had been damaged. We also found that Bt infection led to a decrease in mitochondrial membrane potential, which is well known to be a consequence of mitochondrial ATP (Baker *et al*, 2014). Together, our work revealed Bt infection leads to mitochondria disruption in intestinal cells, increasing the rate of AMP/ATP and cytosolic energy imbalance, which triggers AMPK via AAK-2 phosphorylation. Our work highlighted the importance of the connection between the cellular mitochondria surveillance systems and AMPK through energy changes during pathogen infection.

Additionally, how mitochondria disorder triggering downstream innate immunity signaling is largely unknown yet. Accumulating evidences showed that AMPK is also an important trigger of FOXO family transcription factors. For example, AMPK phosphorylates the transcription factor DAF-16 to increase the transcription of DAF-16-dependent targets in *C. elegans* induced by dietary restriction (Greer *et al*., 2007). DAF-16 regulated by AMPK pathway is also related to defense response against nutrition deprivation and functions to extend lifespan (Fukuyama *et al*., 2012; Greer *et al*., 2007). Here, we found that the AMPK activation mediated by mitochondrial damage regulates the DAF-16 to increase the transcription of certain DAF-16-dependent innate immune effectors, which helps *C. elegans* defend against pathogen infection (Fig. 6). The different sensitivity in several worms against Bt infection indicating that the AMPK is not the only way to activation of DAF-16 (Fig. S13A). At the same time, AMPK may regulate other DAF-16-independent innate immune pathways during Bt infection.

Moreover, the mitogen-activated protein kinases (MAPK) pathway is evolutionarily conserved innate immune pathways (Porta *et al*, 2011; Tan & Shapira, 2011), which has been reported to defend several pathogens infection, including nematicidal Bt and its Cry toxin in *C. elegans* (Griffitts & Aroian, 2005; Huffman *et al*., 2004). MAPK pathway is also reported to be a downstream component of the AMPK signaling pathway in mammal cells (Li *et al*, 2005). The xenophagy for defense against pathogens infection was observably activated by the AMPK-dependent p38 MAPK pathway (Ji *et al*, 2009; Ouimet *et al*, 2016). Under the infection of nematicidal Bt, the up-regulated of 3 typical genes of p38 MAPK pathway (*pmk-1, tir-1* and *sek-1*) together with the activation of PMK-1 demonstrated that AMPK also triggered the p38-MAPK innate immune pathways apart from DAF-16 during Bt infection (Fig. S13B and S13C).

In conclusion, our works demonstrated that host cells can directly sense mitochondrial-mediated intracellular energy imbalance to monitor pathogens, and activate downstream innate immune responses through activation of the AMPK pathway. As modeled in Fig. 7, nematicidal Bt infection in *C. elegans* led to the mitochondria damage and in the energy imbalance of intestinal epithelial cells, the latter of which subsequently triggered AMPK *via* phosphorylation of AAK-2. Then, the AMPK modulates DAF-16-dependent and p38-MAPK-dependent innate immune pathway to defend against nematicidal Bt infection. These findings revealed that AMPK senses the changing of cellular energy imbalance and triggers the innate immune responses during pathogen infection. Consider that the mitochondria surveillance systems and AMPK are conserved components from worms to mammals, our study suggests that disrupting mitochondrial homeostasis to activate the immune system through AMPK-dependent pathways may widely existing in animals, which may provide new strategies for immunotherapy of multiple diseases.

**Figure 7.**
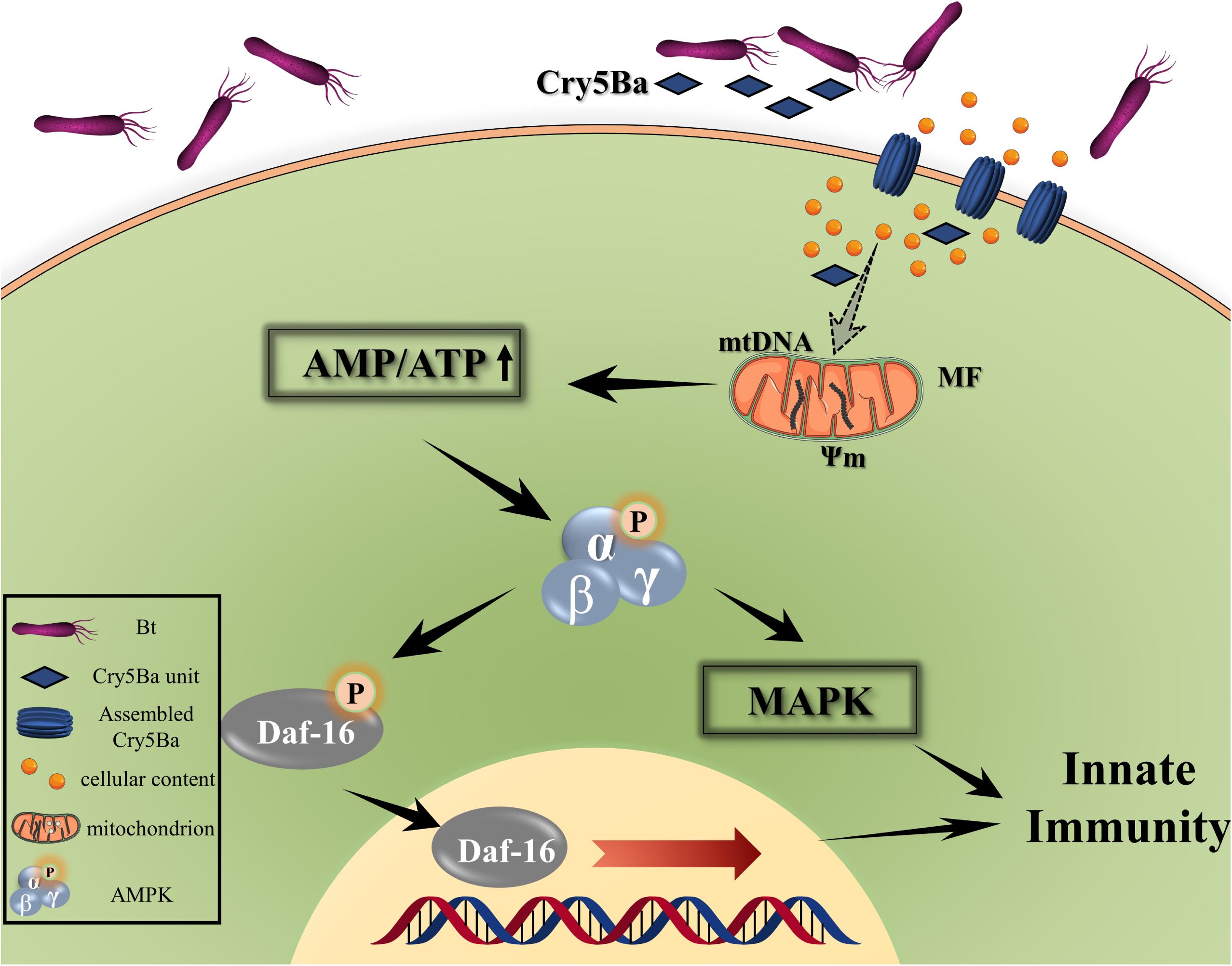
The action model of mitochondria surveillance systems recognition the pathogens and activation innate immune responses by AMPK in *C. elegans*. After nematicidal Bt spores and crystals mixture was fed to the worms, the Cry5Ba toxin assembled to trigger pore formation and transport into the intestinal epithelial cells of *C. elegans*, and disrupts mitochondria indirectly that lead to MF, a decrease in mitochondrial membrane potential, and the changes in mtDNA content, resulting in intercellular energy imbalance. The AMPK senses Bt-caused intercellular ATP changing and activated the AMPK by phosphorylating AAK-2. The activated AAK-2 modulates conversationally DAF-16-dependent and p38-MAPK dependent innate immune pathways to defense pathogens. Solid arrows represent relationship has been confirmed in this study. The dotted arrow represents the relationship is still unclear.

## Materials and Method

### *Caenorhabditis elegans* and bacterial strains

*Caenorhabditis elegans* strains used in this work were kindly provided by the *Caenorhabditis* Genetics Center (CGC) or the National Bioresource Project (NBRP) and listed in Table S1. All *C. elegans* strains were maintained on nematode growth media (NGM, 0.3% NaCl, 0.25% tryptone, and 1.5% agar) using *E. coli* OP50 as food under standard conditions (Brenner, 1974). The bacterial strains and plasmids used in this study are also listed in Table S1. All *Escherichia coli* and Bt strains were grown on Luria-Bertani (LB) medium supplemented with the appropriate antibiotics at 37 °C or 28 °C, for *E. coli* or Bt, respectively. BMB171/Cry5Ba, BMB171/Cry5Ca, BMB171/Cry21Aa, BMB171/Cry6Aa and BMB171/Cry1Ac used in this work is the recombinant Bt that is the acrystalliferous Bt mutant BMB171 transformed with toxin gene *cry5Ba, cry21Aa, cry6Aa* and *cry1Ac* on shuttle vector pHT304 (Guo *et al*, 2008; Peng *et al*, 2009), respectively. BMB171/pHT304 used in this work is the acrystalliferous Bt mutant BMB171 transformed with an empty shuttle vector pHT304.

### Construction of gene *aak-2* rescued worms

A 2.1 kb fragment of the intestine-specific *vha-6* promoter (P_vha-6_), a 2.5 kb fragment of the muscle-specific *myo-3* promoter (P_myo-3_), a 1.3 Kb fragment of the neuron-specific *rab-3* promoter fragment (P_rab-3_) and a 3.0 Kb fragment of *aak-2* promoter fragment (P_*aak-2*_) were generated by PCR from total DNA of *C. elegans*, then inserted into pPD49.26 to gain 4 recombinant plasmids of pPD49.26-P. A full-length a 1.8 Kb of the *aak-2* cDNA and a 1.6Kb of the 3-UTR of *aak-2* was generated by overlap PCR and inserted into pPD49.26-P at the downstream of the promoters respectively. The recombinant plasmids which contained gene *aak-2* with the tissue-specific promoters, P_*aak-2*_::*aak-2*, P_vha-6_::*aak-2*, P_myo-3_::*aak-2* and P_rab-3_::*aak-2*, were injected into the gonads of *aak-2(ok524)* worms to generate independent transgenic lines by standard germline transformation techniques (Merritt & Seydoux, 2010). All of the 4 fusion genes were injected at concentrations ranging 100 ng/μl with P_lin-44_::GFP at 100 ng/μl as a co-injection marker.

### RNA interference

RNAi *E. coli* strains containing targeting genes were originally delivered from the Ahringer RNAi library (Kamath & Ahringer, 2003) and were kindly provided by Professor Zhenxing Wu, Huazhong University of Science and Technology. RNAi feeding experiments were performed as described before (Kamath & Ahringer, 2003) on synchronized L1 larvae at 20°C for 40 h.

### Quantitative real-time RT-PCR analysis

Total RNA isolated using Trizol reagent (Invitrogen) was reversely transcribed with random primers using Superscript II reverse transcriptase (Invitrogen) according to the manufacturer’s protocol. Real-time PCR analysis was performed with Life Technologies ViiA™ 7 Real-Time PCR system (Life Technologies, USA) using Power SYBR Green PCR Master Mix (Life Technologies, USA). The gene *tba-1* was used as an internal reference (Zhang *et al*, 2012). The primers used for PCR are listed in Table S1. The experiments were conducted in triplicate, and the results were expressed as 2^-ΔΔ Ct^.

### Nucleotide measurements

The AMP, ADP and ATP were extracted by the adapting perchloric acid method (Stocchi *et al*, 1985), 300-400 worms were washed with ice-cold M9 buffer, and resuspended in 20 μl of M9 buffer, then added 40µL of ice-cold 8% (v/v) HClO_4_ on ice. The worms were then homogenized in liquid nitrogen, and the homogenates were disrupted using ultrasonic vibrations and neutralized with 1 N KOH. Next, the suspension was centrifuged (10,000g for 3min at 4°C), and the supernatant was passed through a 0.2 µm filter (Nanosep). The concentration of AMP, ADP and ATP were detected using the LC-MS (Agilent Technologies, 1260-6540). The operation of detection is based on the published method(Cortese *et al*, 2019; Wang *et al*, 2014).

### Analyses of mitochondrial morphology

SJ4143(*zcIs17*[P_*ges-1*_::GFP^mt^]) worms can stably express GFP in mitochondria matrix of intestinal cells. It can be used as a reporter for mitochondrial morphology. The worms were synchronized according to the method described before (Urano *et al*., 2002). Then the worms were treated according to the liquid assay. Washed worms were collected to be photographed by Olympus BX63 with excitation at 488 nm and emission at 525 nm. At least three repeats were performed for each condition with and at least 50 animals photographed per treatment.

### Measurement mitochondrial membrane potential (ΔΨm)

Tetramethyl rhodamine ethyl ester (TMRE) ((MCE, # 115532-52-0)) is a lipophilic cation to detect the membrane potential (ΔΨm) of the mitochondrial (Palikaras *et al*, 2015). Synchronized N2 worms were treated according to the liquid assay. After treated by respective bacteria and compounds, worms were resuspended in M9 containing 8μM TMRE. After two hours’ treatment, worms were washed with M9 for 3 times to remove dye thoroughly. Worms were photographed using Olympus BX63 epifluorescence microscope at 10 × magnification with excitation at 550 nm and emission at 575 nm. The average optical density of each worm was measured using Image Pro Plus v6.0. At least three repeats were performed for each condition and at least 30 animals were measured per treatment.

### Mitochondrial DNA (mtDNA) quantification assay

MtDNA damage was assessed using Real Time-qPCR according to the published method. (Cristina *et al*, 2009; Lakshmanan *et al*, 2018) The pair of primers (Forward: GTTTATGCTGCTGTAGCGTG, Reverse: CTGTTAAAGCAAGTGGACGAG) were used to normalize the mitochondrial genome. The results were normalized to genomic DNA using primer pairs specific for *ama-1*(Forward: TGGAACTCTGGAGTCACACC, Reverse: CATCCTCCTTCATTGAACGG). Synchronized worms were treated and collected as described above. Collecting worms were homogenized in liquid nitrogen and lysed in a standard buffer containing proteinase K for1 hour at 65°C. qPCR procedure was implemented as described above.

### Bt infection assay

The Bt infection assays were based on the published protocol (Bischof *et al*., 2006). which contains two different procedures.

### The plates assays

Bt strains were grown in LB medium containing 50μg/ml spectinomycin at 28 °C overnight and seeded 400μl to fresh NGM plate. Then these plates were placed at 28 °C for 3 days. Synchronized nematodes were grown to L4 stage in NGM plate containing OP50. Nematodes were then washed off and cleaned by M9 buffer. 2000-3000 worms were added to the Bt plate and treated at 25°C for 4-6h.

### The liquid assay

Synchronized L1 worms were grown to L4 stage on OP50 plate at 20°C. Then 300-500 worms were added on a 24-well plate (total volume 50 μl) which containing 400μl M9 buffer. Then add different mixture of crystal toxins and spores to each well (total volume 50ul). The plates were placed at 25°C for 4-6h.

### The growth assays

The L1 growth assays were carried out in different doses of the mixture of crystal Cry5Ba and spores as described before (Bischof *et al*., 2006), *E. coli* OP50 was added at an optical density of 0.2–0.25 OD_600_ and 20–30 synchronized L1 larvae were used per well. After 3 days at 20°C, photographed at least 60 worms in different doses on a microscope, and calculated the average area for each condition using the software NIH Image J 1.33, normalized the average area at each toxin concentration to the average area of the no toxin control. The size of the worms in the absence of toxin was set at 100%. Each experiment was independently replicated at least three times.

### The survival assays

The survival assays were conducted in different doses of the mixture of crystal Cry5Ba and spores as described before (Bischof *et al*., 2006). *C. elegans* N2 and these mutant worms were exposed to the mixture of crystal Cry5Ba and spores in S media in 96-well plates to quantitatively score the survival. Concentrations of each toxin were set-up in triplicate for each assay with approximately 20-30 synchronized L4 worms per well. The determination of whether worms were alive is according to the worms’ movement. The crawling worms were marked as alive. Non-crawling worms should be gently touched with a platinum pick to observe their movement. The survival rate of each well was scored after incubating at 20°C for 6 days. Each experiment was independently replicated at least three times.

### Lifespan assays

The life span assay was performed at 20°C on NGM plates. Each bacterium must be fully spread on the NGM plate to prevent worms from avoiding or escaping the bacteria lawn. Approximately 60 L4-stage worms were incubated on NGM plate seeded with OP50 or other pathogens. Every plate contained 0.05 mg/ml of 5-fluorodeoxyuridine (FUDR) to prevent eggs from hatching. Five plates were tested for each strain in each experiment. Each experiment was independently replicated at least three times. The determination of whether worms were alive was as described for the survival assay of *C. elegans*. The surviving worms on each plate were counted at 20 °C every 12 h. Statistical analyses were assessed by Kaplan-Meier survival analysis followed by a log rank test. Statistical significance was assessed by Kaplan-Meier survival analysis followed by a log rank test.

### CuSO_4_ and H_2_O_2_ assay

CuSO_4_ and H_2_O_2_ assay were conducted as previously described (Los *et al*, 2013). The survival assays were conducted in serial doses of CuSO_4_ or H_2_O_2_, 20-30 L4 synchronized worms were used per well in 48-well plates, *E. coli* OP50 at an optical density OD600 was 0.2-0.25. The survival rate of each well was determined after 6 days of CuSO_4_ or 4 hours of H_2_O_2_ exposure at 20°C.

### Cry5Ba localization assay

The L4-staged transgenic worms RT311[P_*vha-6*_::*GFP::RAB-11*], which was an apical recycling endosome reporter (Jiu *et al*, 2012) and SJ4143(*zcIs17* [P_*ges-1*_::GFP^mt^]), which was chosen as mitochondria reporter worm and can stably expressed GFP in mitochondria matrix of intestinal cells(Urano *et al*., 2002), were fed with rhodamine-labeled crystal protein Cry5Ba (a pore-forming toxin produced by Bt) 4 hours, then these worms were placed on 2% agarose pads, the signals of rhodamine-labeled and GFP-labelled were observed using the using confocal microscope at 100×magnification. At least 2-3 independent biological repeats were carried out for each experiment.

### DAF-16 nuclear localization assay

After 2 h of treatment with Bt, the synchronized L4 TJ356 worms (transgenic animals expressing DAF-16::GFP) were immediately placed in M9 buffer and onto microscope slides. GFP localization was observed using a fluorescent microscope (Olympus BX31, Japan) at 40×magnification. DAF-16 localization was categorized as cytosolic localization, intermediate localization and nuclear localization. The number of worms with each level of nuclear translocation was counted. The worms were exposed to non-nematicidal Bt BMB171/pH304 were used as negative controls; while the worms were exposed to heat shock for the same periods at 30°C were used as a positive control. *P* values were calculated using SPSS ver13.0 (SPSS, Chicago, IL).

### Western Blotting Analysis

After feeding Nematicidal *Bt* or Non-nematicidal Bt, *C. elegans* N2 and different mutants were washed three times with ice-cold M9 buffer and were homogenized in liquid nitrogen. Then homogenates were harvested in lysis buffer (50 mM HEPES, 150 mM NaCl, 10% glycerol, 1% Triton X-100, 1.5 mM MgCl2, and 1 mM EGTA) with protein inhibitors (0.2mM Na_3_VO_4_, 1mM NaF). The lysate samples were subjected to SDS-PAGE using 10% (wt/vol) polyacrylamide gels and transferred to PVDF membranes (Life technologies). The trans blotted membrane was washed three times with PBS containing 0.05% Tween 20 (PBST). After blocking with PBST containing 5% BSA, the membrane was probed with the primary antibody (Phospho-AMPKα(Thr172) Rabbit mAb, Cell Signaling, # 2535, or β-actin Antibody, Proteintech, # 66009) and washed three times. The membrane was then probed with HRP-coupled secondary antibody and washed. Finally, the membranes were exposed using a chemiluminescent substrate (SuperSignal West Pico, Thermo Scientific).

### RNA-sequencing and transcriptome analysis

Total RNA was isolated using Trizol reagent (Invitrogen, USA) according to the manufacturer’s protocol. Total RNA concentrations and integrity were measured with NanoDrop 2000C spectrophotometer (Thermo Scientific, USA). The mRNA of each sample was purified using poly-dT oligo attached magnetic beads, and then fragmented (approximately 200 bp) at an increased temperature. The first strand cDNA was synthesized with random oligonucleotides and SuperScript II (Invitrogen, USA). The second-strand cDNA was synthesized by DNA polymerase I and RNase H. The second-strand cDNA was purified with Vahtstm DNA Clean Beads. Ends were repaired and 3-end single nucleotide A (adenine) was added. All RNA-seq libraries (non-strand-specific, paired end) were prepared with the TruSeq RNA Sample Prep kit (Illumina, USA). The 150 nt of sequence was determined from both ends of each cDNA fragment using the HiSeq platform (Illumina) according to the manufacturer. After sequencing, clean reads were obtained by removing low-quality, adaptor-polluted and high content of unknown base (N) reads. Then the clean reads were used to perform de-novo assembly with Trinity (https://github.com/trinityrnaseq/trinityrnaseq/wiki), a TGICL (http://sourceforge.net/projects/tgicl/files/tgicl%20v2.1/) was used to further assemble all the unigenes from the two different groups to form a single set of non-redundant unigenes (called all-unigenes). Sequencing reads were annotated and aligned to the *C. elegans* reference genome using Tophat2. The alignment files from TopHat2 were used to generate read counts for each gene. GO enrichment analysis was implemented by mapping each differentially expressed gene into the records of the GO database (http://www.geneontology.org/). GO terms with corrected p-values less than 0.05 were considered to be significantly enriched by differentially expressed genes. Pathway enrichment analysis was based on the KEGG database. We studied the biologically complex responses of genes and obtained pathway annotation for unigenes with KEGG annotation. A Q-value ≤0.05 was identified as significant enrichment of a pathway among the differentially expressed genes.

### Data Analysis

All data analysis was performed using SPSS, ver20.0 (SPSS, Chicago, IL, USA). Statistical analysis between two values was compared with a paired *t*-test. Statistical analysis among three or more values was compared with one-way ANOVA with Dunnett adjustment. Statistics indicated are: *: *p*< 0.05, **: *p*< 0.01, ***: *p*< 0.001. The lack of any symbol indicates no significant difference.

## Data accessibility

mRNA-sequencing data are available on the NCBI Sequence Read Archive (SRA) (https://www.ncbi.nlm.nih.gov/sra), under the bioproject PRJNA662857 (https://dataview.ncbi.nlm.nih.gov/object/PRJNA662857?reviewer=dnhoor4a4e7johdlqepcm0d53h).

## Acknowledgement

This work was supported by the fund from National Key R&D Program of China [2017YFD0200400 to D.P.]; National Natural Science Foundation of China [31670085 to M.S. and 31770116 to D.P.]; China 948 Program of Ministry of Agriculture [2016-X21 to M.S.]; National Natural Science Foundation of China [31600006 to S.J.]; China Postdoctoral Science Foundation [2016M602316 to S.J.]; the Natural Science Foundation of Hubei province [2018CFB529 to S.J.]; and the College Excellent Youth Science and Technology Innovation Team Project of Hubei Province [T201535 to S.J.]. We thank Professor ZhengXing Wu for providing RNAi strains (College of Life Science and Technology, Huazhong University of Science and Technology). We also thank the Caenorhabditis Genetics Center for the worm strain. Microscopic image data were acquired at the State Key Laboratory of Agricultural Microbiology Core Facility.

## Author contributions

Shouyong Ju conceived the project, Shouyong Ju and Hanqiao Chen designed and performed experiments. Shaoying Wang and Jian Lin assist to performed the experiments. Shouyong Ju and Hanqiao Chen complete the writing of the manuscript. Donghai Peng and Ming Sun designed the experiments and revised the paper.

